# Adgrg6/Gpr126 is required for myocardial Notch activity and N-cadherin localization to attain trabecular identity

**DOI:** 10.1101/2022.05.27.493401

**Authors:** Swati Srivastava, Felix Gunawan, Alessandra Gentile, Sarah C. Petersen, Didier Y.R. Stainier, Felix B. Engel

## Abstract

How adhesion G protein-coupled receptors (aGPCRs) control development remains unclear. The aGPCR Adgrg6/Gpr126 has been associated with heart trabeculation. Defects in this process cause cardiomyopathies and embryonic lethality. Yet, how cardiomyocytes attain trabecular identity is poorly understood. Here, we show that Gpr126 regulates Notch activity and N-cadherin localization that are necessary for attaining trabecular identity in zebrafish. Maternal zygotic *gpr126^stl47^* early truncation mutants exhibit hypotrabeculation whereby N-cadherin distributes randomly at apical/basal membranes of compact layer cardiomyocytes. In contrast, *gpr126^st49^* mutants expressing a N-terminal fragment lacking the GPS motif (NTF^ΔGPS^) exhibit a multilayered ventricular wall consisting of polarized cardiomyocytes with normal N-cadherin expression and increased Notch signaling. Notably, endocardially expressed C-terminal fragment (CTF) reinstates trabeculation in *gpr126^st49^* mutants. Collectively, our data indicate domain-specific roles of Gpr126 during trabeculation whereby the NTF maintains cell-cell adhesion and is required for compact wall integrity, while the CTF is essential to provide trabecular identity

## Introduction

Adhesion G protein-coupled receptors (aGPCRs) are required for the regulation of various biological and developmental processes^1^. Concordantly, mutations in aGPCRs have been associated with a wide variety of human diseases^1–3^. Moreover, ∼34% of all FDA-approved drugs target 108 members of the GPCR superfamily^4^. Therefore, uncovering the function of understudied GPCRs provides a wealth of untapped therapeutic potential. aGPCRs are characterized by a seven-transmembrane (7TM) domain and a long extracellular domain (ECD) consisting of various adhesion domains. They are autoproteolytically cleaved within a conserved GPCR Autoproteolysis INducing (GAIN) domain into a C-terminal fragment (CTF) containing the 7TM domain and an N-terminal fragment (NTF) containing the ECD^5, 6^. The signaling mechanisms of aGPCRs and their domain specific functions are still poorly understood. Various models of receptor activation have been proposed^7–9^ while there are also reports that aGPCRs use their ECDs to mediate functions in a 7TM-independent manner^10^.

Adgrg6/Gpr126 is essential for the development of several tissues/organs including the peripheral nervous system, intervertebral disc, ear, placenta, and heart^11–15^. Several genome-wide Association (GWAS) and RNAseq studies have associated variants in the human *GPR126* locus with diseases like adolescent idiopathic scoliosis (AIS), intellectual disability, severe arthrogryposis multiplex congenita, periodontitis, chronic obstructive pulmonary disease (COPD), and various human cancers^16–18^. In regard to heart development, it has been demonstrated that global deletion of *Gpr126* affects cardiac trabeculation in mice^13, 15, 19^ and zebrafish^13^. Waller-Evans *et al.* described the first *Gpr126* knock out mouse line, which was characterized by mid-gestation lethality exhibiting circulatory failure like congestion, edema, and internal hemorrhage with secondary defects in trabeculation and a normally developed placenta^19^. Subsequently, two different *Gpr126* knockout mouse models were reported exhibiting embryonic lethality at E11.5, which is associated with hypotrabeculation, myocardial wall thinning, arrhythmia, bradycardia, and mitochondrial defects^13, 20^. Morpholino-mediated *gpr126* knockdown in zebrafish also resulted in hypotrabeculation and rescue experiments using Gpr126-NTF suggested that the Gpr126-NTF is sufficient for heart development^13^. In Gpr126 knockout mice, however, it has been proposed recently that placental defects account for the heart abnormalities^15^.

Cardiac trabeculation is a crucial process during ventricular chamber development where the cardiomyocytes in the heart wall organize themselves to form two distinct layers, an outer compact wall and an inner trabecular wall. This process depends on communications between the epicardium, myocardium, and endocardium^21, 22^. Endocardial Notch signaling is a critical early regulator of trabeculation^23^. Subsequently, Neuregulin (Nrg1 in mice /Nrg2a in zebrafish) is cleaved and secreted from the endocardium and induces ErbB2 signaling in the myocardium^24–30^. Cardiomyocyte crowding triggers their delamination, activating Notch signaling in a subset of compact wall cardiomyocytes^31–33^. Myocardial Notch activation inhibits compact wall cardiomyocyte delamination by suppressing the actomyosin machinery, preserving myocardial wall architecture^33^. Notch-negative cardiomyocytes depolarize and delaminate, adopting a trabecular identity. During trabeculation, the adherens junctions between cardiomyocytes are remodeled. N-cadherin, the major component of cardiac adherens junctions, moves in compact wall cardiomyocytes from the lateral to the basal domain. In contrast, trabecular cardiomyocytes exhibit a punctate N-cadherin distribution on their entire surface^34^. The genetic network underlying this Notch-mediated fate specification remains to be elucidated.

In this study, we utilized genetic zebrafish models with domain-specific loss and gain of function approaches coupled with high resolution microscopy to assess the role of Gpr126 in heart development. Our data suggest that both the NTF and CTF of Gpr126 are required for trabeculation. The NTF of Gpr126 contributes to maintaining compact wall integrity at the onset of trabeculation by maintaining cell-cell junctions. The CTF helps in providing trabecular identity to cardiomyocytes through modulation of myocardial Notch activity. Altogether our results provide new insights into the domain-specific functions of Gpr126 and shed light on the regulation of the mechanisms underlying Notch-mediated fate specification during trabeculation.

## Results

### The *stl47* and *st49* mutant alleles produce truncated versions of Gpr126

In order to understand the role of Gpr126 as well as its NTF and CTF, the zebrafish *gpr126* mutant lines *gpr126^stl47^* and *gpr126^st49^* were analyzed. The *stl47* allele carries a 5+3 insertion deletion (indel) at Gln68 which produces a frameshift-induced stop codon in the CUB domain, resulting in a truncated Gpr126 protein of 96 amino acids (Figure S1A, B). The *st49* allele carries a point mutation that converts a tyrosine-encoding codon into a stop codon near the GPS motif; as such, *gpr126^st49^* mutant alleles could encode a truncated Gpr126 protein of 782 amino acids corresponding to the NTF without the GPS motif (NTF^ΔGPS^) (Figure S1A, B). Both mutations result in a puffy ear phenotype caused by abnormal semicircular canal morphogenesis (Figure S1C)^35^. qPCR analysis revealed no differences in *gpr126* expression levels in hearts obtained from 96 hpf wild type (WT), *gpr126^stl4^*^7^, and *gpr126^st4^*^9^ embryos (Figure S1D). These data suggest that the mutant *gpr126* mRNAs are not subjected to nonsense-mediated RNA decay.

Thus far, it has not been tested whether the predicted truncated proteins are produced from the *gpr126^stl47^*and *gpr126^st4^*^9^ alleles. As no working anti-Gpr126 antibodies are currently available, we ectopically expressed zebrafish full length *gpr126* cDNA, containing the *stl47* or *st49* mutation as well as an N-terminal HA-tag and a C-terminal GFP-tag (Figure 1A), in the human retinal pigment epithelial cell line ARPE-19, which is routinely used for protein expression studies. Immunofluorescence analysis showed that WT as well as mutant forms of Gpr126 were expressed (Figure 1B). Notably, for the mutant forms, although the N-terminal HA signal was present, no C-terminal GFP signal was detected, indicating that the mutations resulted in truncated forms of Gpr126 (Figure1B, C). WT Gpr126 localized to the plasma membrane 24 hours post transfection (Figure 1B). In contrast, the mutant Gpr126 forms failed to localize to the membrane and were both detected in the cytoplasm, which is in accordance with loss of the 7TM domain in both truncated forms. To support further that the predicted truncated Gpr126 forms are expressed, western blot analyses were performed to determine the size of the encoded proteins. Anti-GFP antibodies detected a band of ∼70 kDa in the WT-transfected cells which corresponds to Gpr126-CTF-GFP (69 kDa) (Figure 1D). No bands were detected in cells expressing the mutant forms, confirming the immunofluorescence results (Figure 1B, C). Anti-HA antibodies detected two bands of ∼90 and ∼100 kDa in the WT-transfected cells which correspond to HA-NTF^ΔGPS^ (91 kDa) (Figure 1D). Similarly, a band of ∼90 kDa was detected in cells expressing the *st49*-mutant form, which is predicted to encode the NTF^ΔGPS^. In cells expressing the *stl47*-mutant form, a band of ∼10 kDa was detected, the size of which corresponds to the predicted truncated Gpr126 protein containing the signal peptide and the partial CUB domain (11 kDa).

**Figure 1.**
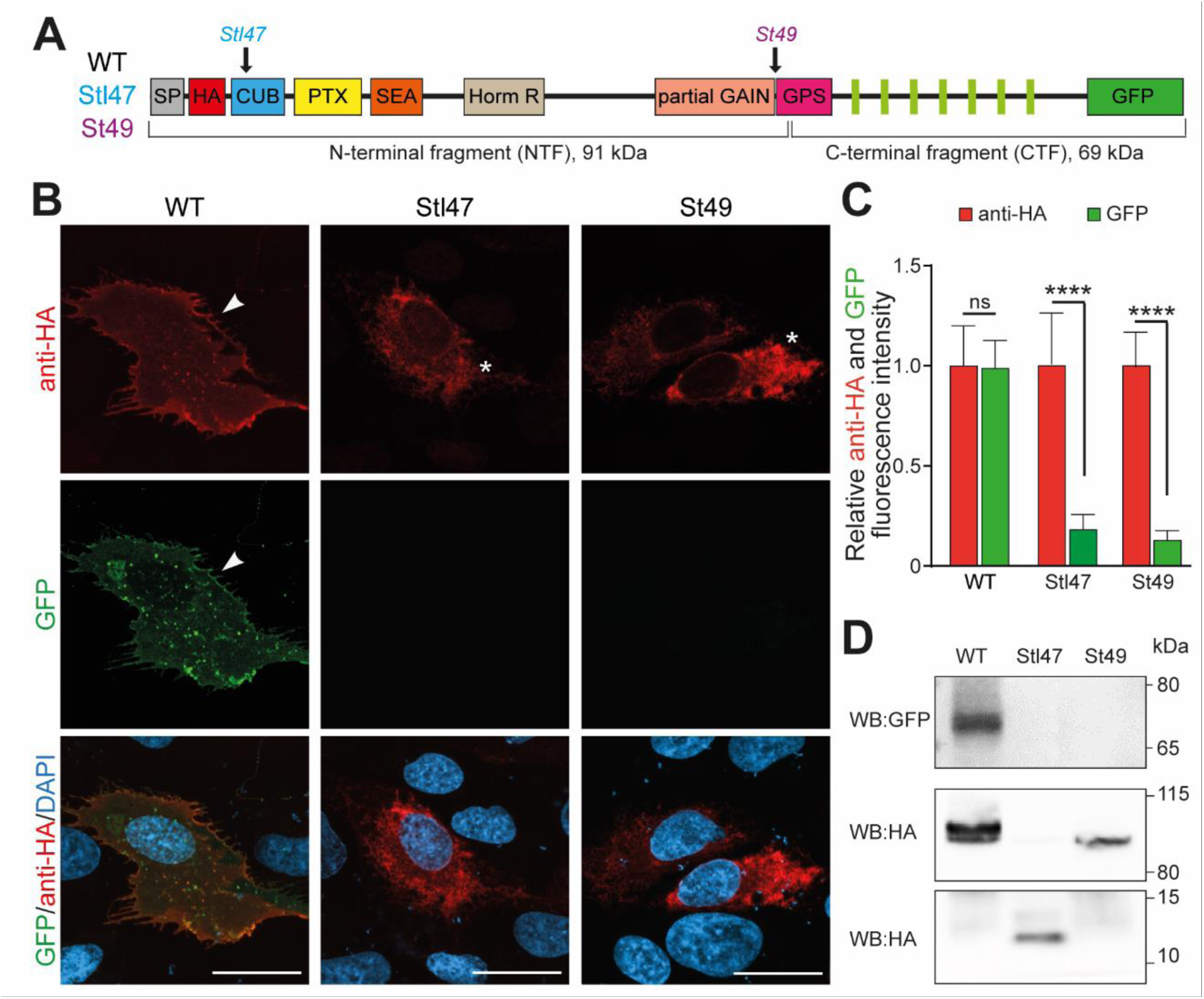
*gpr126* mutants produce truncated Gpr126 proteins. **(A)** Schematic representation of Gpr126 protein tagged with HA at its N-terminus and GFP at its C-terminus indicating *stl47* and *st49* mutation sites. **(B)** Representative images of ARPE-19 cells transfected with *HA-gpr126-GFP* (WT), *HA-gpr126^stl47^-GFP* (stl47), and *HA-gpr126^st49^-GFP* (st49) stained for HA. Nuclei were visualized with DAPI. Arrowheads: membrane localization of HA and GFP. Asterisks: intracellular localization of HA. Scale bars: 10 µm. **(C)** Quantification of (B). Fluorescence intensity was normalized to intensities of *HA-gpr126-GFP* (WT)-transfected cells. Data are mean ± SD. ****: p< 0.0001 by one-way ANOVA. ns: not significant. **(D)** Representative Western blot (WB) analysis of lysates from ARPE-19 cells transfected with *HA-gpr126-GFP* (WT), *HA-gpr126^stl47^-GFP* (stl47), and *HA-gpr126^st49^-GFP* (st49) using antibodies against GFP and HA; as indicated.

Altogether, these data demonstrate that the expression of zebrafish full length *gpr126* cDNA containing the *stl47* or *st49* mutation in ARPE-19 cells results in the expression of the predicted truncated forms of Gpr126 and indicate that the zebrafish mutant *gpr126^stl47^* expresses the first 98 amino acids of Gpr126 containing a part of the CUB domain while *gpr126^st49^* expresses NTF^ΔGPS^. Consequently, analysis of these two mutant zebrafish lines will provide insight into the role of Gpr126 and its individual NTF and CTF domains.

### Maternal zygotic *gpr126^stl47^* mutants exhibit hypotrabeculation

Previous reports aiming to define the role of Gpr126 during heart development are conflicting^13,15,19^. In order to determine whether Gpr126 is required for heart development, the *gpr126^stl47^* line, predicted to express only the first 98 amino acids of Gpr126, was crossed with the reporter line *Tg(myl7:EGFP-hsa.HRAS)* in which GFP is localized at the cardiomyocyte plasma membrane. *In vivo* confocal imaging of the hearts of WT siblings showed clear luminal projections into the ventricle forming a complex network of trabeculae at 96 and 120 hpf (Figure 2A). The analysis of zygotic *gpr126^stl47^* mutant hearts revealed no obvious gross morphological defects compared to WT siblings suggesting that Gpr126 is not required for heart development in zebrafish (Figure S2). However, it has previously been reported that *gpr126^stl47^* maternal zygotic (MZ) mutants, which are the progeny of homozygous mutant females, exhibit a more severe phenotype during Schwann cell development. Unlike zygotic mutants, MZ mutants have no maternal deposition of wildtype mRNA or protein into the egg and showed more pronounced defects in *mbp* expression than zygotic mutants^35^. Similarly, we observed trabeculation defects in MZ *gpr126^stl47^* mutants at 96 and 120 hpf. In 16.7% of MZ *gpr126^stl47^* mutant larvae, we observed that trabeculation was absent and the ventricular wall was single-layered; furthermore, in 26.7% of MZ *gpr126^stl47^* mutant larvae, hypotrabeculation was detected, and 20% of the hearts showed an abnormal multilayered ventricular wall (Figure 2A, B). Thus, a total of 63.4% of the MZ *gpr126^stl47^* mutants exhibited a trabeculation defect. These data suggest that maternal contribution of Gpr126 rescues the zygotic loss in *gpr126^stl47^* mutants and that Gpr126 is required for cardiac chamber development in zebrafish.

**Figure 2.**
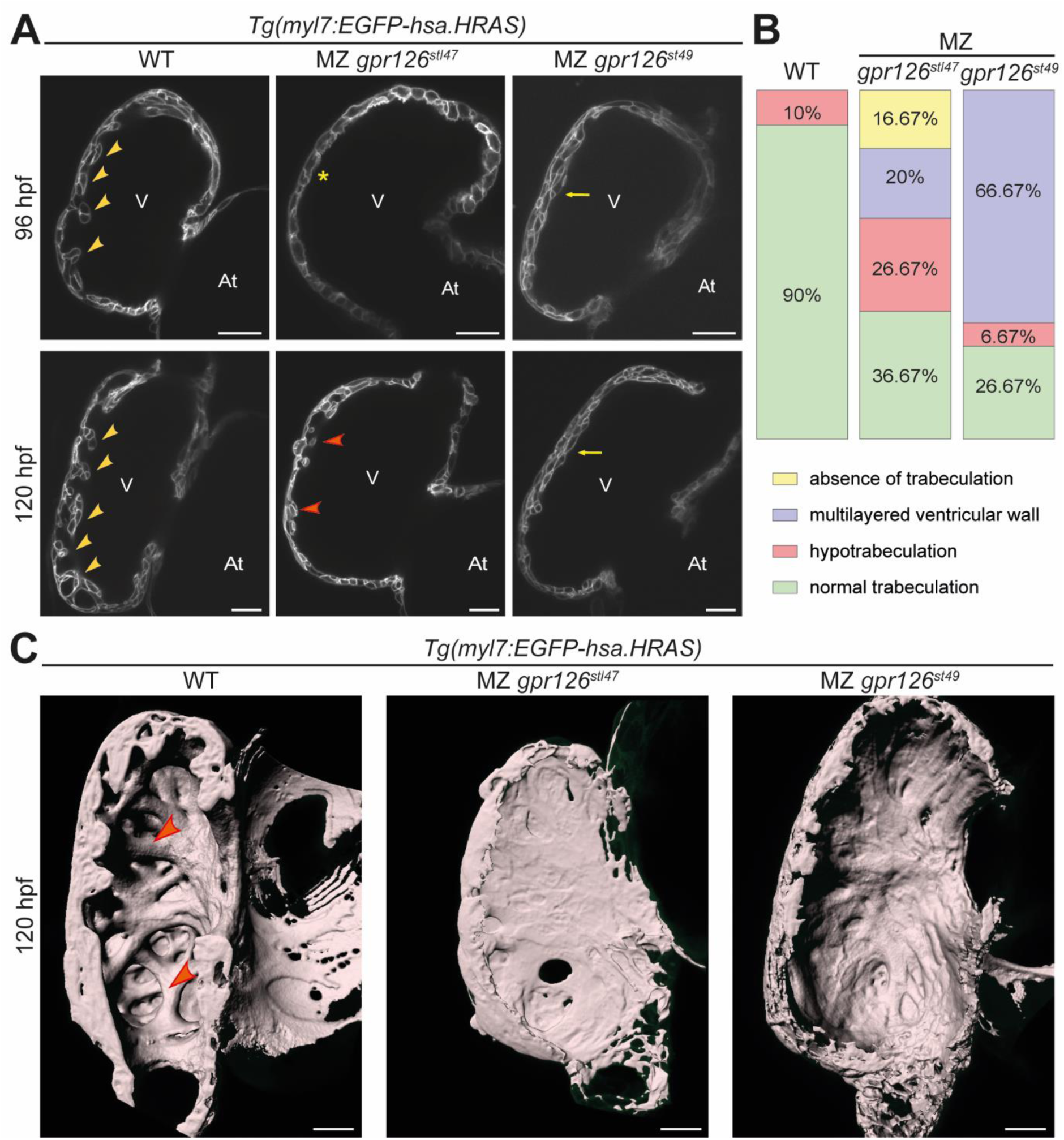
Gpr126 NTF and CTF are required for trabeculation. **(A)** Representative confocal images (mid-sagittal sections) of hearts of WT, MZ *gpr126^stl47^*, and MZ *gpr126^st49^* larvae crossed into *Tg(myl7:EGFP-hsa.HRAS)* background at 96 and 120 hpf. Yellow arrowheads: trabeculae, orange arrowheads: shorter trabeculae Asterisks: absence of trabeculae. Arrows: multilayered ventricular wall. V: ventricle. At: atrium. Scale bars: 20 µm. **(B)** Quantification of trabeculation phenotypes in 120 hpf WT (n=20), MZ *gpr126^stl47^* (n=30), and MZ *gpr126^st49^* (n=30) larvae. MZ: maternal zygotic. **(C)** 3D surface rendering of the myocardium of 120 hpf WT and mutants as indicated. Arrowheads: trabecular projections. Scale bars: 20 µm.

### Gpr126-CTF is required for cardiac trabeculation

Previously, morpholino-mediated knockdown experiments suggested that the NTF^ΔGPS^ is sufficient for trabeculation and heart development^13^. To further test this model, considering the issues associated with morpholino studies^36, 37^, we crossed the *gpr126^st49^* line, predicted to express only NTF^ΔGPS^, with the reporter line *Tg(myl7:EGFP-hsa.HRAS)*. We first analyzed zygotic mutants and observed that the majority of zygotic *gpr126^st49/st49^* hearts exhibit a multilayered ventricular wall (60%) and absence of trabecular ridges (Figure S2). The MZ *gpr126^st49^* mutants recapitulate the multilayered phenotype (66.67%) observed in the zygotic mutants (Figure 2A, B). In addition, 6.67% of the mutants exhibited hypotrabeculation. These data indicate that the CTF of Gpr126 is required for proper cardiac trabeculation.

For further characterization of the phenotypes of *gpr126^stl47^* and *gpr126^st49^* mutants, we analyzed MZ animals and their WT cousins, as control, by performing IMARIS-based 3D surface reconstruction of the ventricular chambers at 120 hpf. In WT larvae, distinct muscular ridges in the ventricular lumen could be visualized. In contrast, MZ *gpr126^stl47^* mutants exhibited a smooth ventricular wall with no trabecular projections, while MZ *gpr126^st49^* mutant hearts exhibited a thick compact wall with few or no luminal protrusions (Figure 2C).

Collectively, these data indicate that Gpr126 is required for heart development in zebrafish whereby the different phenotypes of *gpr126^stl47^* and *gpr126^st49^* mutants suggest that both NTF and CTF are required for trabeculation but play different roles in this morphological process.

### Endocardial Gpr126-CTF expression reinstates trabeculation in *gpr126^st49^* mutants

In mouse, it has been demonstrated that *Gpr126* is an endocardially expressed gene^13, 15^. We performed fluorescence activated cell sorting (FACS) for endocardial cells and cardiomyocytes using isolated cells from extracted hearts of *Tg(myl7:EGFP-hsa.HRAS)*; *Tg(kdrl:nls-mCherry)* larvae at 48 and 96 hpf. Endocardial and myocardial enrichment was confirmed by the relative expression levels of *kdrl* and *myl7*, respectively. qPCR analysis confirmed that in zebrafish hearts, *gpr126* is also enriched in endocardial cells compared to cardiomyocytes (Figure 3A). Therefore, in order to perform rescue experiments in *gpr126^st49^* mutants, we generated a stable endothelial-specific transgenic line, *Tg(fli1a:gpr126 CTF-p2a-tdTomato)* that overexpresses Gpr126 CTF-p2a-tdTomato, which is post-translationally cleaved into CTF and tdTomato. *Tg(fli1a:gpr126 CTF-p2a-tdTomato)* larvae did not show any gross morphological defects (Figure S3A) and were raised to adulthood. The analysis of 96 hpf *Tg(fli1a:gpr126 CTF-p2a-tdTomato)* larval hearts showed normal trabeculation and no obvious heart defects (Figure S3B).

**Figure 3.**
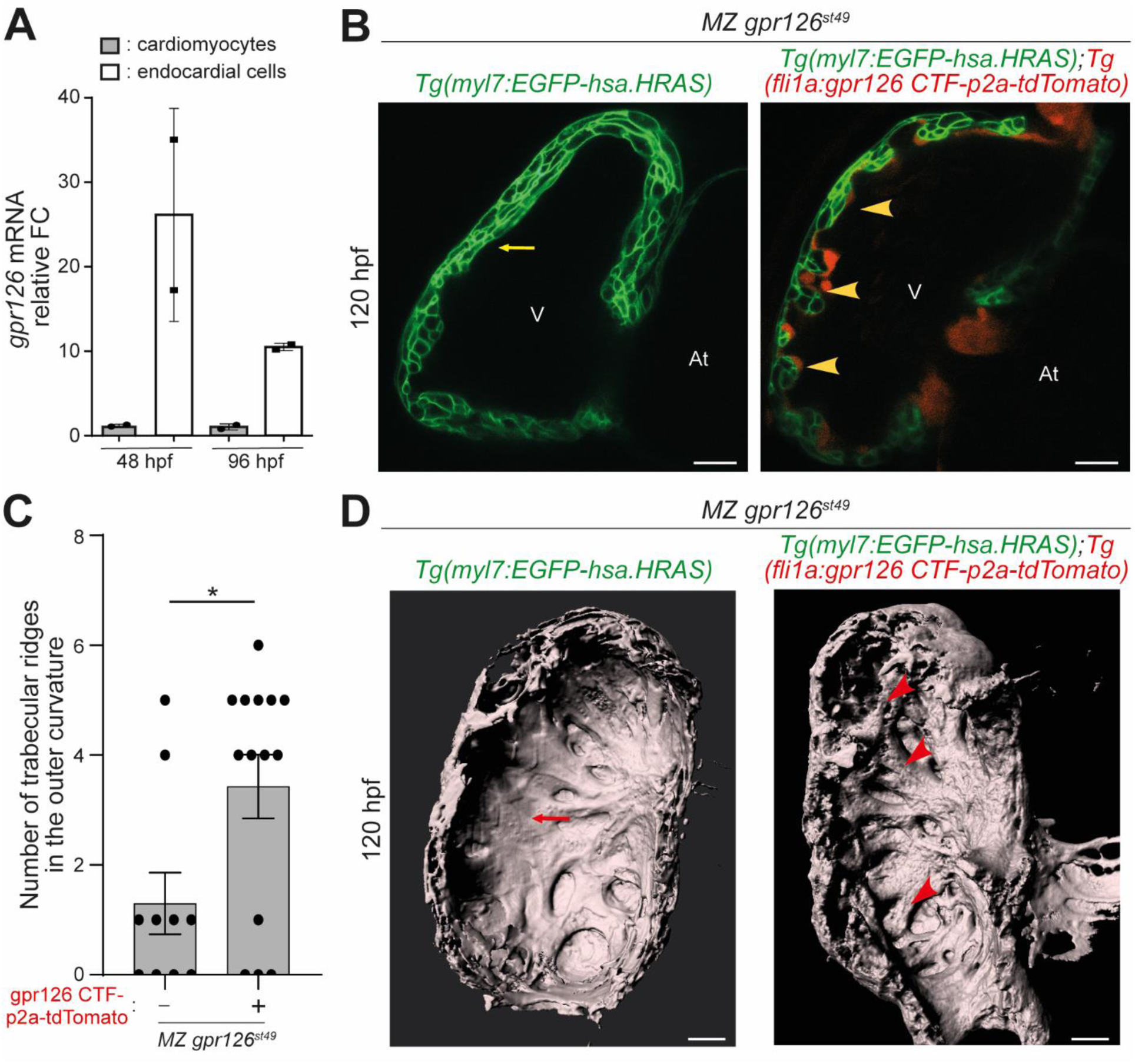
Endocardial CTF overexpression reinstates trabeculation in MZ *gpr126^st49^* mutants. **(A)** qPCR data from FACS sorted cardiomyocytes and endocardial cells from hearts at 48 and 96 hpf. Relative mRNA levels of *gpr126* are enhanced in endocardial cells compared with the cardiomyocytes. n=2 biological replicates. FC: fold change. Data are mean ± S.E.M. **(B)** Representative confocal images (mid-sagittal sections) of *Tg(my7:EGFP-has.HRAS);* MZ *gpr126^st49^* and *Tg(my7:EGFP-has.HRAS);Tg(fli1a:gpr126 CTF-p2a-tdTomato);* MZ *gpr126^st49^* hearts at 120 hpf. Yellow arrows: multi-layered ventricular wall; yellow arrowheads: trabeculae; V: Ventricle; At: Atrium. Scale bars: 20 µm. **(C)** Quantification of (B). Number of trabecular ridges in MZ *gpr126^st49^* hearts in the absence (n=8) or presence (n=14) of Gpr126 CTF-P2A-TdTomato. Dots represent individual hearts. Data are mean ± S.E.M. *: p= 0.0183 by Student’s t-test. **(D)** 3D surface rendering of the myocardium of 120 hpf Tg(my7:EGFP-has.HRAS); MZ *gpr126^st49^* and *Tg(my7:EGFP-has.HRAS);Tg(fli1a:gpr126 CTF-p2a-tdTomato)*; MZ *gpr126^st49^* hearts. Red arrowheads: trabecular projections; red arrows: thick compact wall with absence of mature trabecular ridges. Scale bars: 20 µm.

To determine whether CTF overexpression in the endocardium could rescue the multilayered ventricular wall in MZ *gpr126^st49^* mutants, *Tg(fli1a:gpr126 CTF-p2a-tdTomato)* were crossed with *gpr126^st49^* mutants in the background of *Tg(myl7:EGFP-hsa.HRAS)* and raised to adulthood. The analysis of 120 hpf MZ *gpr126^st49^; Tg(fli1a:gpr126 CTF-p2a-tdTomato)*; *Tg(myl7:EGFP-hsa.HRAS*) hearts revealed that ectopic CTF expression, identified by tdTomato-positive endocardium, reversed the multilayering phenotype in MZ *gpr126^st49^* mutant hearts, resulting in the formation of trabecular projections into the ventricular lumen (Figure 3B, C). This was confirmed by 3D surface rendering of the myocardium. MZ *gpr126^st49^* mutants overexpressing CTF exhibited clear muscular ridges in the ventricular chamber, while MZ *gpr126^st49^* mutants exhibited a thick compact wall in the absence of mature trabecular ridges (Figure 3D), as described above (Figure 2C). These data indicate that endocardial expression of Gpr126-CTF rescues the multilayering phenotype observed in *gpr126^st49^* mutants.

Collectively, our data suggest that endocardial-specific Gpr126-CTF signaling is required for proper trabeculation in zebrafish.

### MZ Gpr126*^st49^* mutant and Gpr126 overexpression phenotypes are ErbB2-dependent

In order to determine the effect of full length Gpr126 overexpression on trabeculation, we generated the transgenic line *Tg(fli1a:gpr126-p2a-tdTomato)* to overexpress Gpr126 in all endothelial cells including the endocardium. The F2 generation of *Tg(fli1a:gpr126-p2a-tdTomato);Tg(myl7:EGFP-hsa.HRAS)* exhibited normal gross morphology and survived to adulthood. However, confocal microscopy analysis of 120 hpf *Tg(fli1a:gpr126-p2a-tdTomato)* hearts revealed that the ventricular wall is multilayered (7/8) (Figure 4A). In addition, these hearts exhibited a novel ventricular aneurysm^38, 39^-like phenotype (5/8) characterized by pockets in the ventricular wall, internally lined by endocardium (Figure 4A).

**Figure 4.**
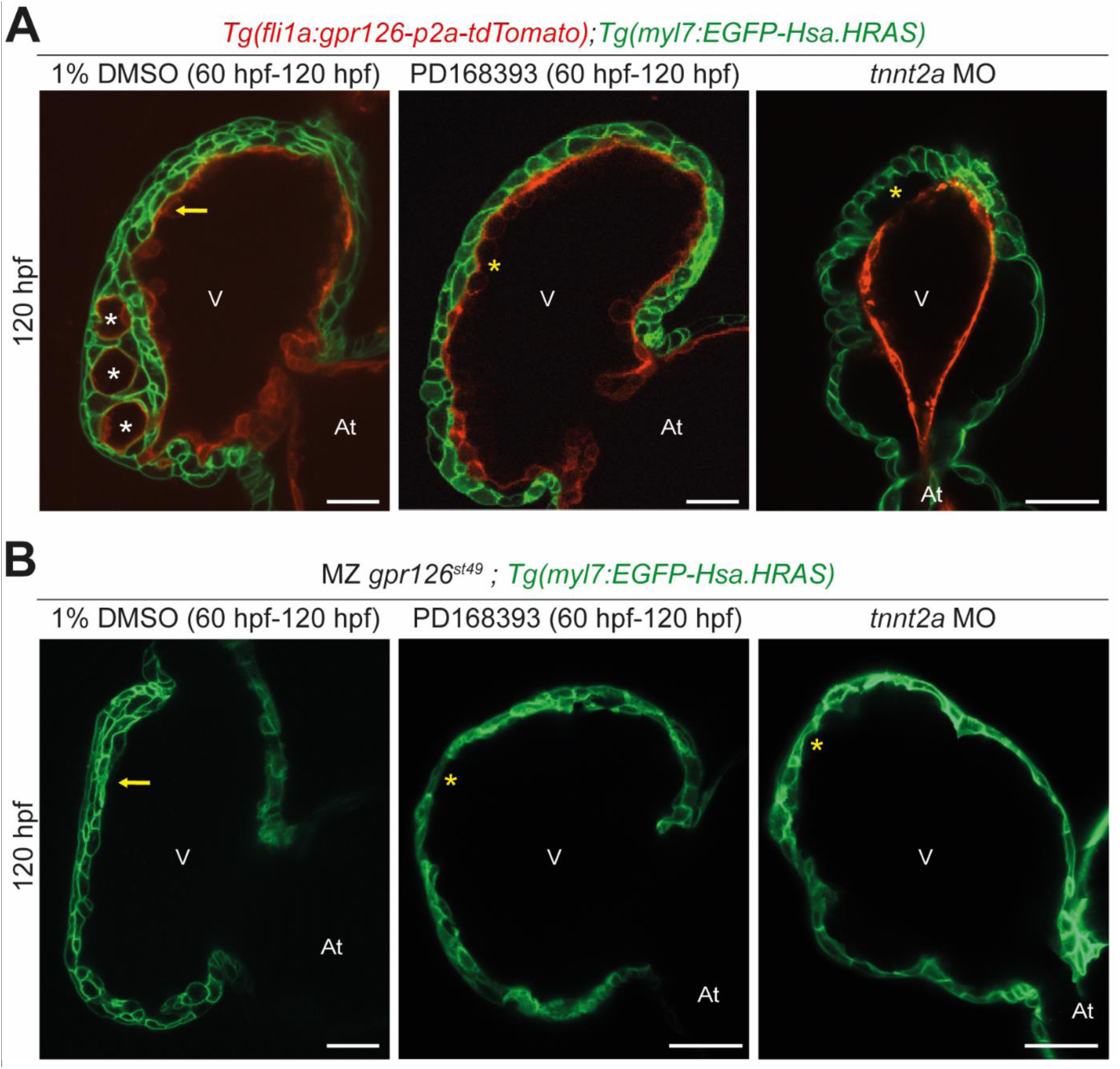
ErbB2 signaling and blood flow and/- contractility are required for gpr126 function. **(A)** Confocal images (mid-sagittal sections) of 120 hpf *Tg(fli1a:gpr126-p2a-tdTomato);Tg(my7:EGFP-hsa.HRAS)* hearts treated with 1% DMSO or PD168393 or injected with *tnnt2a* morpholino. White asterisks: aneurysm; yellow arrows: multilayered ventricular wall; yellow asterisks: absence of trabeculae; V: Ventricle; At: Atrium. Scale bars: 20 µm **(B)** Confocal images (mid-sagittal sections) of 120 hpf MZ *gpr126^st49^;Tg(my7:EGFP-hsa.HRAS)* hearts treated with 1% DMSO or PD168393 or injected with *tnnt2a* morpholino. Yellow arrows: multilayered ventricular wall; yellow asterisks: absence of trabeculae; V: Ventricle; At: Atrium. Scale bars: 20 µm

ErbB2 signaling as well as blood flow and/or contractility are crucial regulators of trabecular morphogenesis and lack of either of these factors abrogates trabeculation^26, 27^. To assess whether Gpr126 signaling is dependent or independent of ErbB2 signaling, the major signaling pathway known to control trabeculation^26, 27, 29, 30^, we treated *Tg(fli1a:gpr126-p2a-tdTomato);Tg(myl7:EGFP-hsa.HRAS)*and *gpr126^st49^; Tg(myl7:EGFP-hsa.HRAS)* larvae with 1% DMSO or 10 μM of the ErbB2 inhibitor PD168393 from 60 to 120 hpf. In both animal models, ErbB2 inhibition blocked trabeculation resulting in hearts with single-layered ventricular walls (Figure 4A, B). These data suggest that ErbB2 signaling is required for the action of Gpr126. Similarly, inhibition of blood flow and cardiac contractility utilizing *tnnt2a* morpholinos, resulted at 120 hpf in both animal models in a single-layered heart (Figure 4A, B).

Taken together, these data suggest that the regulation of trabeculation via Gpr126 signaling depends on ErbB2 signaling and blood flow or contractility.

### Gpr126-CTF is required to establish trabecular identity of cardiomyocytes

The compact layer cardiomyocytes exhibit apicobasal polarity, while trabecular layer cardiomyocytes are depolarized^32^. The observed multilayering phenotype of *gpr126^st49^* MZ mutants raised the question whether the inner layer of cardiomyocytes consist of depolarized cardiomyocytes, indicating hypertrabeculation, or whether they consist of polarized cardiomyocytes caused by abnormal multilayering of the compact ventricular wall. In order to determine the apicobasal polarity of cardiomyocytes, MZ *gpr126^st49^* mutants were crossed into the background of *Tg(myl7:EGFP-Podxl);Tg(myl7:mKATE-CAAX)*, in which the cardiomyocyte membrane is labeled in red with mKATE and the apical membrane of polarized cardiomyocytes is marked by the apical protein podocalyxin fused with EGFP. In WT siblings at 96 hpf, EGFP-podocalyxin is localized at the apical/abluminal side of compact layer cardiomyocytes and around the entire membrane of delaminated, trabecular cardiomyocytes, indicating their polarized and depolarized states, respectively (Figure 5A). In contrast, in multilayered hearts of MZ *gpr126^st49^* animals, cardiomyocytes of both the outer layer as well as the inner layers were mainly polarized (Figure 5A). Quantification revealed that the percentage of depolarized cardiomyocytes in the inner layers of the ventricles was significantly reduced from 94.7% ± 1.7% in WT to 16.6% ± 2.6% in MZ *gpr126^st49^* mutants (p < 0.0001; WT: n = 6 hearts; MZ *gpr126^st49^*: n = 12 hearts; per heart 15-25 cardiomyocytes were counted) (Figure 5B). These data indicate that the cardiomyocytes in multilayered hearts of MZ *gpr126^st49^* mutants fail to depolarize and consequently cannot delaminate to seed the trabecular layer.

**Figure 5.**
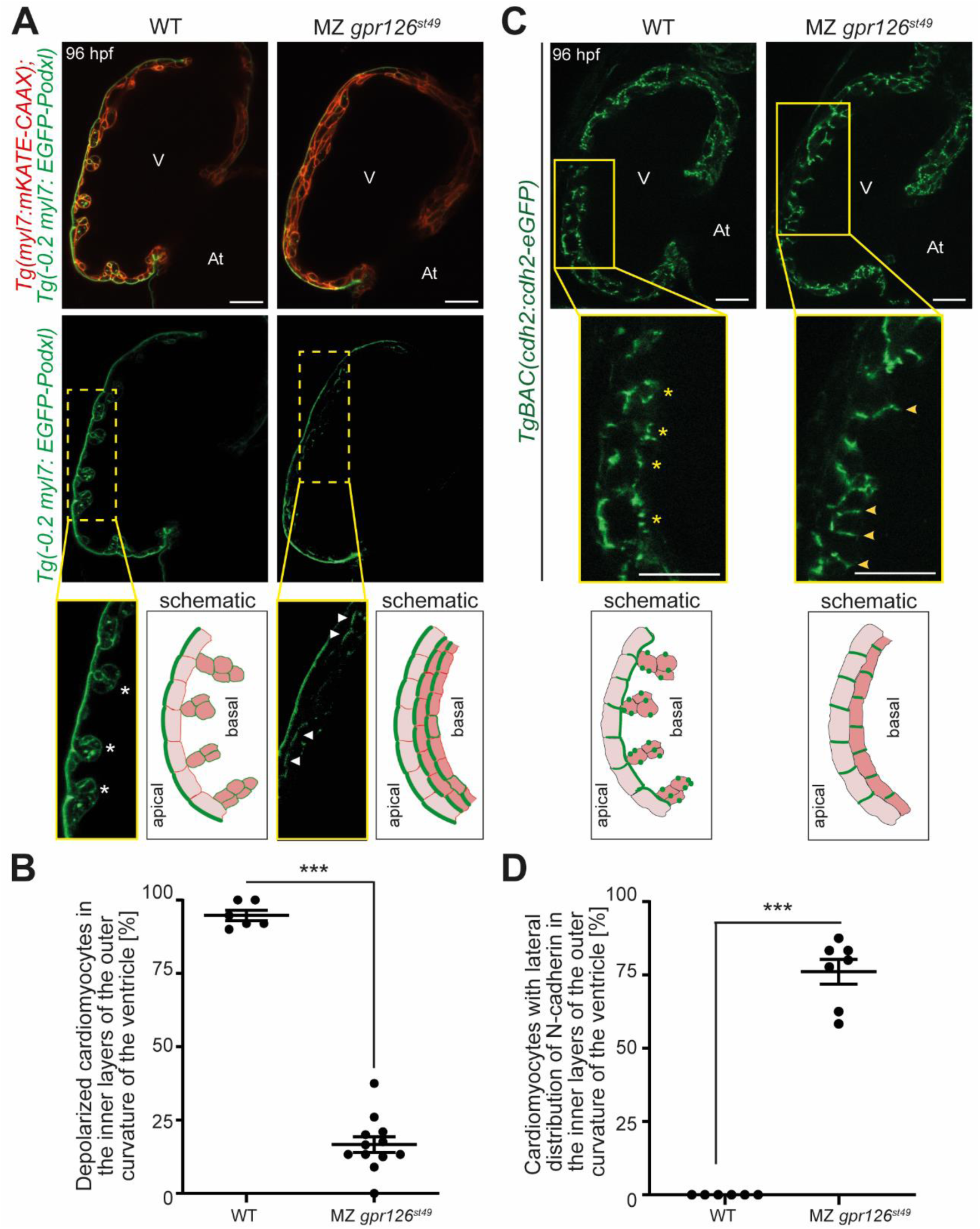
Gpr126 CTF aids in providing trabecular identity to cardiomyocytes. **(A)** Representative confocal images (mid-sagittal sections) of 96 hpf *Tg(myl7:EGFP-Podxl); Tg(myl7:mKATE-CAAX);* WT and MZ *gpr126^st49^* hearts. Asterisks: depolarized and delaminated cardiomyocytes; arrowheads: polarized cardiomyocytes; V: ventricle; At: atrium. Scale bars: 20 µm. Schematic represents the depolarized cardiomyocytes in the trabecular layer of WT hearts and polarized cardiomyocytes in the inner layers of *gpr126^st49^* MZ hearts at 96 hpf. **(B)** Percentage of depolarized cardiomyocytes in the inner layers of the outer curvature of the ventricles of WT (n = 6) and MZ *gpr126^st49^* (n = 12) larvae. Each point represents a heart. Data are mean ± S.E.M. ***: p< 0.001 by Student’s t-test. **(C)** Representative confocal images (mid-sagittal sections) of 96 hpf *TgBAC(cdh2:cdh2-EGFP);* WT (10/10) and MZ *gpr126^st49^* (11/14) hearts. Asterisks: punctate distribution of N-cadherin around the cardiomyocyte; arrowheads: lateral distribution of N-cadherin; V: Ventricle; At: Atrium; Scale bars: 20 µm. Schematic represents N-cadherin localization in WT and MZ *gpr126^st49^* hearts at 96 hpf. **(D)** Percentage of cardiomyocytes with lateral localization of N-cadherin in the inner layers of the outer curvature of the ventricles of WT (n = 6) and MZ *gpr126^st49^* (n = 7) larvae. Each point represents a heart. Data are mean ± S.E.M. ***: p< 0.001 by Student’s t-test.

During depolarization, compact layer cardiomyocytes remodel their adherens junctions, which involves the relocalization of N-cadherin from the lateral to the basal membrane. When the trabecular cardiomyocytes leave the compact layer, they exhibit punctate N-cadherin distribution on the surface facing the basal side of compact layer cardiomyocytes^34^. To test whether adherens junction remodeling is affected in MZ *gpr126^st49^* mutant hearts, N-cadherin localization was analyzed at 96 hpf, when multilayering was observed, utilizing the reporter line *TgBAC(cdh2:cdh2-EGFP)*, where N-cadherin (Cdh2) is labeled with EGFP. In WT hearts, the compact layer cardiomyocytes showed basolateral distribution of N-cadherin and trabecular layer cardiomyocytes had a punctate distribution. In contrast, N-cadherin in MZ *gpr126^st49^* mutant hearts was localized at the lateral side of the cardiomyocytes in the outer as well as inner multilayered wall (Figure 5C, D).

Collectively, our data indicate that cardiomyocytes in the multilayered wall of *gpr126^st49^* mutants fail to attain trabecular identity due to errors in remodeling adherens junctions and cardiomyocyte polarity defects.

### Gpr126-NTF is sufficient to maintain cell-cell adhesion and compact wall integrity

To determine if NTF^ΔGPS^ is sufficient to maintain proper N-cadherin localization in compact wall cardiomyocytes prior to trabeculation, *gpr126^stl47^* and *gpr126^st49^* mutants crossed in the background of *TgBAC(cdh2:cdh2-EGFP)* were analyzed at 72 hpf. Single-plane confocal images of WT hearts showed that N-cadherin localizes laterally in the membrane of compact layer cardiomyocytes (Figure 6). Maximum intensity projection images confirmed the junctional localization of N-cadherin in WT hearts. In contrast, in MZ *gpr126^stl47^* mutant hearts, N-cadherin was distributed randomly at the apical or basal membrane of the compact layer cardiomyocytes, which was also obvious in maximum intensity projections. Notably, in the MZ *gpr126^st49^* mutants, which express NTF^ΔGPS^, N-cadherin localization was, similar to the WT siblings, distributed at the junctions of the compact layer cardiomyocytes. These data suggest that the NTF^ΔGPS^ is sufficient for the proper localization of adherens junctions in cardiomyocytes of the compact wall prior to trabeculation.

**Figure 6.**
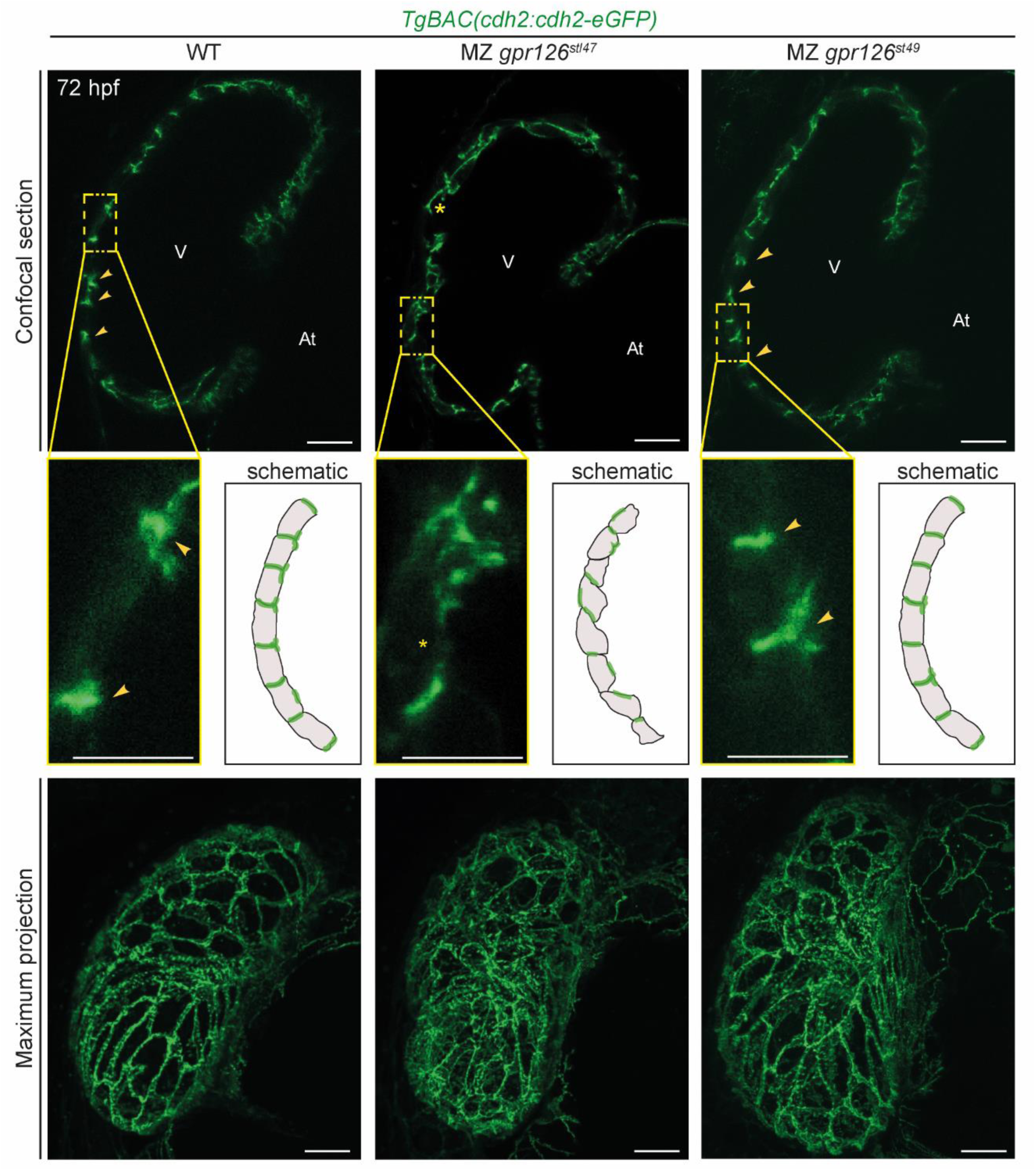
Gpr126 NTF regulates compact wall integrity at the onset of trabeculation. Representative confocal images (mid-sagittal sections and maximum intensity projections) of *TgBAC(cdh2:cdh2-GFP);* WT (10/10) as well as MZ *gpr126^stl47^* (10/12) and MZ *gpr126^st49^* (12/12) hearts at 72 hpf. Arrowheads: lateral localization of N-cadherin; asterisks: mislocalization of N-cadherin; V: ventricle; At: atrium. Scale bars: 20 µm, 10 µm (magnified images).Schematic represents N-cadherin localization in the compact layer of WT, MZ *gpr126^stl47^* and MZ *gpr126^st49^* hearts at 72 hpf.

To further investigate the independent role of NTF, Gpr126 NTF^ΔGPS^ was overexpressed in a mosaic manner by injecting single cell-stage *Tg(myl7:EGFP-hsa.HRAS)* embryos with a *myl7:gpr126NTF-p2a-tdTomato* plasmid. NTF overexpressing cardiomyocytes were identified by tdTomato expression. Confocal microscopy analysis revealed that NTF overexpression in cardiomyocytes leads to local multilayering at 96 hpf in 62% of injected hearts (5/8). The tdTomato-overexpressing cardiomyocytes adhere together along with tdTomato-negative cardiomyocytes, while the rest of the ventricular wall is monolayered (Figure S4). This result suggests that NTF overexpression increases cell adhesion resulting in the formation of a cardiomyocyte multilayer, providing further evidence that NTF has a function in maintaining cell-cell adhesion in the compact layer cardiomyocytes.

### Gpr126-CTF modulates myocardial Notch activity to provide trabecular identity

It has been suggested that Notch signaling is activated in distinct compact layer cardiomyocytes at the onset of trabeculation, which prevents them from becoming trabecular cardiomyocytes^31, 33^. Therefore, we hypothesized that failure of *gpr126^st49^* mutant cardiomyocytes to attain trabecular identity might be due to enhanced myocardial Notch activity. To investigate whether Notch signaling is modulated by CTF or NTF, *gpr126^stl47^* and *gpr126^st49^* mutants were crossed in the background of *Tg(TP1:Venus-PEST)*, in which the Notch-responsive TP1 promoter drives expression of Venus-PEST, a fluorescent protein with a short half-life (∼2 h). This allows to identify cells which have recently activated Notch signaling. During trabeculation, Notch signaling can be detected in the atrioventricular canal (AVC), the outflow tract and a few cardiomyocytes in the compact layer of the myocardium. Analysis of the mutant hearts at 96 and 120 hpf indicated that Notch activity was not altered in MZ *gpr126^stl47^* mutants (Figure 7A). In contrast, Notch activity in the heart was markedly increased in MZ *gpr126^st49^* mutants. Quantitative analysis of the Venus fluorescence intensity normalized to the signal in AVC valves confirmed that myocardial Notch activity was significantly increased in MZ *gpr126^st49^* mutants (Figure 7B). This suggests that CTF signaling is required to inhibit myocardial Notch activity to allow cardiomyocytes to attain trabecular identity.

**Figure 7.**
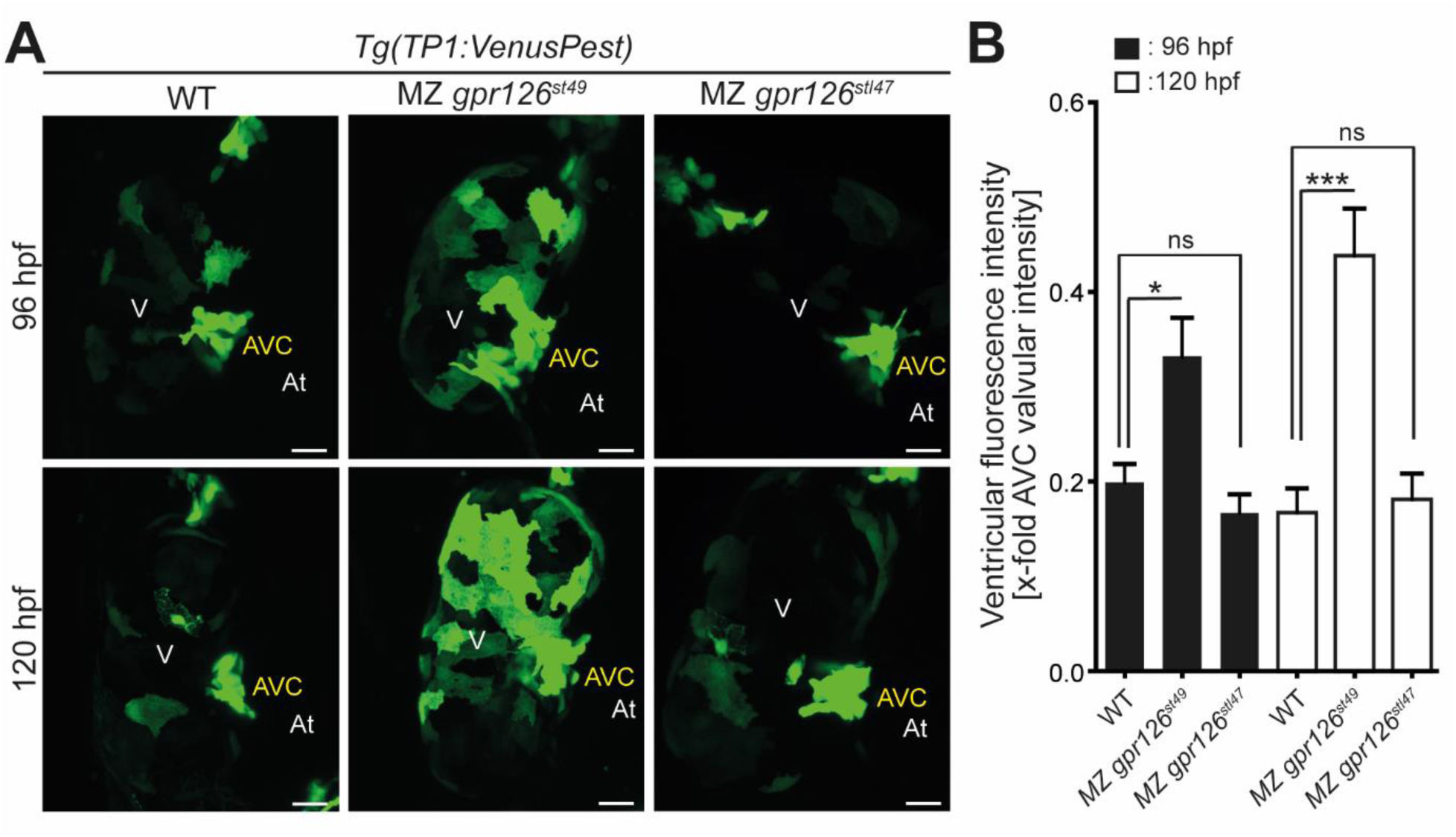
*gpr126* modulates myocardial notch activity. **(A)** Maximum intensity projections of representative *Tg(TP1:Venus-PEST);* WT, MZ *gpr126^stl47^,* and MZ *gpr126^st49^* hearts showing active notch signaling at 96 and 120 hpf. MZ *gpr126^st49^* hearts express significantly higher myocardial Notch signaling activity as compared to WT and MZ *gpr126^stl47^* hearts. Scale bars: 20 µm. **(B)** Quantification of Venus-Pest intensity with reference to valves for WT, MZ *gpr126^stl47^,* and MZ *gpr126^st49^* hearts at 96 hpf (n=7) and 120 hpf (n=8). Data are mean ± S.E.M. *: p< 0.05; ***: p< 0.001; ns: not statistically significant by one way ANOVA with corrections for multiple comparisons by Dunnett’s test.

In MZ *gpr126^st49^* mutants, the NTF is no longer bound to the CTF. Thus, secreted NTF acting on the myocardium might be responsible for the observed MZ *gpr126^st49^* mutant phenotypes. To test this hypothesis, we established the *Tg(myl7:gpr126NTF-p2a-tdTomato)* zebrafish line and analyzed whether ectopically expressed NTF^ΔGPS^ in the myocardium mirrors the *gpr126^st49^* MZ mutant phenotypes. Utilizing *Tg(TP1:Venus-PEST)* revealed that myocardial Notch signaling was not enhanced at 96 hpf (Figure S5). These data suggest that the increased myocardial Notch activity in MZ *gpr126^st49^* mutants is due to impaired CTF signaling in endocardial cells and not due to an increased availability of NTF.

## Discussion

We conclude that Gpr126 is required for heart development. First, MZ *gpr126^stl47^* mutants, which encode a severely truncated Gpr126 protein, exhibit hypotrabeculation. Second, *gpr126^st49^* hearts, with a truncated CTF but an intact NTF, are mainly multilayered, characterized by defects in adherens junction remodeling and lack of depolarization. Third, endocardial Gpr126-CTF expression reinstates trabeculation in *gpr126^st49^* mutants. Fourth, cell-cell adhesion and compact wall integrity is maintained in MZ *gpr126^st49^* mutant hearts. Finally, myocardial Notch signal was markedly increased in MZ *gpr126^st49^* mutant hearts. Thus, our data indicate that both NTF and CTF regulate distinct cellular mechanisms of ventricular chamber development. The NTF is required to maintain cell-cell adhesion in the compact layer in order to maintain compact wall architecture prior to the onset of trabeculation. The CTF aids in providing trabecular identity to cardiomyocytes by remodeling adhesion junctions and modulating myocardial Notch activity.

Previously, it has been reported that loss of Gpr126 results in hypotrabeculation^13^. However, no cellular or molecular mechanisms underlying this phenotype were identified. A recent study suggested that the observed heart defects in Gpr126 knockout mice are secondary to placental defects^15^. Yet, in Gpr126 knockout mouse models^12, 13, 15, 19^, placenta defects were not always observed,^12, 13, 19^ and zebrafish do not have a placenta. Moreover, the lack of heart trabeculation defects in the recently analyzed *gpr126* mutants *gpr126^bns341^* (predicted early truncation) and *gpr126^bns342^* (promoter-less allele) is in agreement with our data showing that zygotic *gpr126^stl47^* mutant hearts exhibit no obvious trabeculation defect^15^. However, congruently with a more severe phenotype during Schwann cell development in MZ *gpr126^stl47^* mutants^35^, we observed hypotrabeculation phenotypes in MZ *gpr126^stl47^* mutants. This suggests that the maternal deposition of mRNA rescues the heart phenotypes in zygotic mutants, including *gpr126^bns341^* and *gpr126^bns342^* mutants. Recently, a mechanism of genetic compensation termed transcriptional adaptation has been described in zebrafish, where mutations that cause mutant mRNA degradation might trigger the upregulation of potential compensatory genes^40, 41^. Thus, it is possible that the absence of trabeculation phenotypes in the *gpr126^bns341^* and *gpr126^bns342^* mutants result from a possible compensation through other adapting genes.

Previously reported rescue experiments in *gpr126* morphants suggested that NTF^ΔGPS^ is sufficient for heart development^13^. Notably, no markers were used to distinguish compact and trabecular layer cardiomyocytes. Furthermore, morpholinos exhibit short-lived effects and the latest analyzed time point was at 80 hpf. Therefore, the observed rescue potential of NTF^ΔGPS^ needed to be validated. Here we show that *gpr126^st49^* mutants, which express NTF^ΔGPS^, exhibit from 96 hpf onwards a multilayered ventricular wall consisting mainly of compact layer cardiomyocytes. This suggests an important role of the CTF in trabeculation, which is further substantiated by the fact that endocardial-specific overexpression of CTF rescued the multilayering phenotype in *gpr126^st49^* mutant hearts. The data presented in this work corrects our previous hypothesis that NTF is sufficient for trabeculation and provides evidence that the CTF has an indispensable role in regulating trabeculation.

The N-terminal regions of aGPCRs consist of various adhesion domains which are capable of mediating cell-cell and cell-matrix interactions. While various studies have confirmed a role of aGPCRs in maintaining cell-cell adhesion *in vitro*^42, 43^, there is limited knowledge *in vivo*. Cardiomyocytes are linked to each other through intercalated discs (ICD) which allow the heart to function as a syncytium, with adherens junctions being a major component of ICDs^44^. The adherens junction defects observed in the cardiomyocytes of MZ *gpr126^stl47^* mutants prior to the emergence of trabeculae reveal an early role of Gpr126 in maintaining cell-cell adhesion in the compact wall. As MZ *gpr126^st49^* mutants do not exhibit these defects, the NTF appears to have an independent role in the proper localization of adherens junctions in the compact layer supporting our previous conclusion that NTF acts as a secreted ligand^13^.

Further analysis of CTF-deleted MZ *gpr126^st49^* mutants revealed the failure of cardiomyocytes to depolarize and modulate their adherens junctions. These results suggest that CTF-deleted cardiomyocytes fail to attain trabecular identity and cannot delaminate to seed the trabecular layer, leading to the formation of a multilayered compact ventricular wall. The dual role of NTF and CTF in ventricular wall morphogenesis is similar to a previous study reporting a dual role for the hippo pathway effector Wwtr1 in maintaining compact wall architecture as well as modulating trabeculation^45^.

Notch-mediated fate specification has emerged as a defining feature of trabecular morphogenesis^31, 33^. However, how myocardial Notch activity is regulated remains elusive. Notch activity in compact layer cardiomyocytes actively suppresses their delamination and prevents them to enter into the trabecular layer^33^. The enhanced myocardial Notch signaling in CTF-deleted MZ *gpr126^st^*^49^ mutants are in line with these studies and explain why the CTF-deleted cardiomyocytes fail to attain trabecular identity and stay in the compact layer. These results suggest a negative regulation of myocardial Notch activity by CTF signaling and thus links Gpr126 with Notch signaling-driven lateral inhibition during trabecular emergence.

While our data demonstrate that endocardially expressed Gpr126 is required for regulating cardiomyocyte behavior, it remains unclear how. Endocardial-myocardial interactions are essential for ventricular chamber development and several endocardial factors like Notch, Nrg2a, and Klf2 have previously been reported to affect myocardial patterning^28, 30, 46^. However, how these factors affect cardiomyocyte behavior remains largely unknown and is currently under investigation. Recently it has been suggested that endocardial protrusions regulates communication between endocardium and myocardium by modulating Nrg/ErbB2 signaling^47, 48^. We favor a model in which Gpr126 either regulates the formation of endocardial protrusions or is transported to the myocardium through these protrusions. Previously, we have provided evidence that NTF is acting as a secreted ligand similar to Nrg2a^15^. Thus, it is possible that the CTF is mainly required to control the release of the NTF from the endocardium to the myocardium. However, ectopic expression of NTF^ΔGPS^ in the myocardium, simulating the situation in the *gpr126^st49^* mutants, did affect cell-cell adhesion but failed to enhance myocardial Notch activity. These data further support that both NTF and CTF are required for myocardial patterning. It is likely that CTF regulates endocardial function through canonical G protein-coupled signaling, as observed in myelination during peripheral nervous system development^7,35^. Our study provides further evidence that aGPCRs play a role in cell adhesion in addition to cellular signal transduction^35, 49^.

Overall, our findings enhance our understanding of how aGPCRs function *in vivo*, clarify the importance of Gpr126 in heart development, and provide new insights into how myocardial Notch and cell-cell adhesion between cardiomyocytes is regulated. This is of great importance considering that aGPCR mutations are associated with human diseases^50–52^, congenital heart disease is the most common congenital disease affecting around 1% of live birth^53–55^, and the therapeutic potential of targeting aGPCRs^56^.

## Methods

### Zebrafish maintenance

All animal experiments were performed according to German Animal Protection Laws approved by the local governmental animal protection committee. Embryonic and adult zebrafish were raised and maintained in Tecniplast aquaculture main and stand-alone systems. The water temperature was maintained at 28.5°C and the pH and conductivity was set at 7-7.5 and 450-500 µS. The light-dark cycle consisted of half an hour of gradual onset of light (8 am to 8:30 am) with 13.5 hours of uninterrupted light and half an hour of gradual onset of darkness with 11.5 hours of complete darkness. Zebrafish were fed three times a day, depending on age, with rotifers and granular food. Health monitoring was performed at least once per year.

### Zebrafish mutant and transgenic lines

In this study, the following transgenic and mutant lines were used: Tg(myl7:EGFP-Hsa.HRAS)s883^57^; Tg(−0.2myl7:EGFP-podocalyxin)bns10^32^; Tg(myl7:mKate-CAAX)sd11, abbreviated as myl7:mKate-CAAX^58^; TgBAC(cdh2:cdh2-eGFP, crybb1:ECFP)zf517, abbreviated as TgBAC(cdh2:cdh2-GFP)^59^; Tg(EPV.Tp1-Mmu.Hbb:Venus-Mmu.Odc1)s940, abbreviated as TP1:VenusPest^60^; gpr126^stl47 35^; gpr126^st49 61^; Tg(fli1:gpr126-p2a-tdTomato); and Tg(fli1:tdTomato-p2a-gpr126CTF).

### Genotyping and matings of *gpr126* mutants

*stl47* were genotyped as previously described^35^. For *st49*, the primers listed in Table S1 were used to amplify a 604-base pair (bp) product from genomic DNA. The point mutation in st49 introduces a *Mae*III recognition site and the restriction digestion of the PCR product yields 437-bp and 167-bp fragments. The 604-bp product is retained in the digested WT DNA. For generating maternal zygotic (MZ) animals, homozygous *gpr126^stl47^* or *gpr126^st49^* mutant females were crossed with their heterozygous male counterparts. The WT siblings from heterozygous crosses were grown to adulthood and their progeny were used as controls for MZ experiments, and were referred to as WT cousins.

### RNA isolation from zebrafish larvae heart and quantitative PCR

Larval zebrafish hearts were isolated from WT and mutant embryos in the background of *Tg(myl7:EGFP-hsa.HRAS)* at 120 hpf following the protocol used by Burns and MacRae^62^. Larvae (n = 200) were first anesthetized using 0.04% tricaine methanesulfonate (MS-222) and collected into 1.5 ml microcentrifuge tubes. The larvae were washed three times with ice-cold embryo disruption medium (EDM) and were resuspended in 1.25 ml EDM. Hearts were then fragmented from the larvae by mechanical disruption using a 5 ml syringe and a 1½ inch long 19-gauge needle. This was done by collecting the contents of the tube in the syringe through the needle and expelling it back in the microcentrifuge tube 30 times at the rate of 1 s per syringe motion. Fragmented larvae were filtered through a 105 μm nylon mesh and the flow-through was collected in a 30 mm petri-dish. The flow-through was then applied to a 40 μm nylon mesh which was then inverted and the retained material was collected with EDM in a 30 mm petri dish. The beating hearts were identified with their EGFP expression under the Zeiss stereoscope and were collected into a 1.5 ml microcentrifuge tube containing RNAlater. All these steps were performed on ice. The hearts were then pelleted for 10 min at 10,000 x g. The supernatant was discarded and TRIzol was added onto the pellet and RNA was isolated using RNeasy Micro Kit (Qiagen) according to manufacturer’s instructions. For production of cDNA, 500 ng of RNA was reverse transcribed using Oligo (dT) 12-18mer primers and M-MLV Reverse Transcriptase (Thermo Fisher Scientific, Cat# 280250139) according to the manufacturer’s instructions. qPCR assays were performed in experimental triplicates using SYBR Green (Bio-Rad) in a CFX Connect™ Real-Time PCR (Bio-Rad). *gpr126* expression values were normalized using the housekeeping gene *rpl13a* and fold changes were calculated using the 2^-ΔΔCt^ method. The primers for *gpr126* amplifies part of exon 3 and 4 and the sequences used for qPCR can be found in Table S1.

### Plasmid constructs

For overexpression of zebrafish *gpr126* and its mutant forms in mammalian cell lines, constructs were created using site-directed mutagenesis. HA-gpr126 fragment was amplified by PCR from pcDps-HA-gpr126-FLAG gifted by Ines Liebscher and cloned in the mammalian expression vector pAcGFP to create HA-*Gpr126*-GFP construct. Inserts with the *stl47* and *st49* mutations were generated through PCR by introducing complementary mutations in the primers, as listed in Table S1. The inserts with the mutations were assembled using the NEBuilder HiFi assembly master mix (New England Biolabs, Cat# E2621L) according to the manufacturer’s instructions, and transformed into competent bacteria. To generate the *fli1a:gpr126-p2a-tdTomato* and *fli1a:gpr126 CTF-p2a-tdTomato* plasmids, sequences encoding the Gpr126 and Gpr126 CTF were PCR amplified. The amplified coding sequences followed by *p2a-tdTomato* were cloned downstream of the endothelial specific *fli1a* promotor into a Tol2 enabled vector using the Cold Fusion Cloning kit (System Biosciences, Cat# MC010B-1).

### ARPE19 cells and plasmid transfection

The ARPE-19 cell line was cultured in Dulbecco’s modified Eagle’s medium/Nutrient mixture F-12 (DMEM/F-12) supplemented with 10% fetal bovine serum (FBS) and 100 U/ml penicillin, and 100 μg/ml streptomycin. Cells were maintained in a humidified atmosphere with 5% CO_2_ at 37°C. Plasmid transfection into ARPE-19 cells was carried out with 500 ng DNA per well of a 24-well plate using 1 µl Lipofectamine LTX (Thermo Fisher Scientific, Cat# 15338100)) according to the manufacturer’s instructions. Transfection complexes were formed by incubating DNA with Lipofectamine LTX in Opti-MEM for 20 min at room temperature. For western blot, transfection was performed in 6-well plates and the volumes were adjusted by using fourfold amounts of reagents.

### Immunofluorescence

For detection of HA-tag via immunofluorescence staining, cells were rinsed once with D-PBS and were fixed with 4% formaldehyde/PBS for 10 min at room temperature. Formaldehyde-fixed cells were permeabilized for 10 min with 0.5% TritonX-100/PBS. Prior to antibody staining, samples were blocked for 20 min using 5% BSA in 0.2% Tween-20 in PBS. The primary antibody (Abcam anti-HA) were diluted (1:500) in the blocking reagent and incubated with the sample overnight at 4°C in a humidified chamber. After removal of primary antibody solution and three 5 min washes with 0.1% NP40/PBS, samples were incubated for 60 min with fluorophore-coupled secondary antibodies diluted in blocking reagent (1:500). DNA was visualized with 0.5 μg/ml DAPI (4′,6′-diamidino-2-phenylindole) in 0.1% NP40/PBS. After DAPI staining, cover slips were rinsed once with Millipore-filtered water and then mounted using Fluoromount-G mounting medium.

### Western blotting

Transfected ARPE19 cells were washed with ice-cold PBS and extracted in RIPA buffer supplemented with EDTA-free protease inhibitor cocktails (cOmplete, Roche # 11873580001). After 30 min incubation on ice, samples were sonicated and lysates were cleared by centrifugation at 16,000 x g for 10 min at 4°C. Lysates were analyzed by SDS-PAGE (4-12% NuPAGE Novex Bis-Tris gels) under reducing conditions and transferred to nitrocellulose membrane by wet transfer at 30 V and 300 mA for 1.5 h in 1x transfer buffer (25 mM Tris-HCl, pH 7.5, 192 mM glycine, 0.1% SDS, 10% methanol). The membrane was then blocked with 1x TBS, 0.05% Tween-20 and 10% non-fat milk and incubated with primary antibodies against HA and GFP. Enhanced chemiluminescence reagent (PerkinElmer, Waltham, MA) was used for protein detection.

### FACS sorting and quantitative PCR

Larval hearts (*Tg(myl7:GFP-CAAX)*;*Tg(kdrl:nls-mCherry)*) at 48 and 96 hpf were manually dissected in DMEM+10% FBS, centrifuged for 3 min at 5000 rpm, washed in 1 ml of Hanks’ Balanced Salt Solution (HBSS), and dissociated into single cells in 100 μl of Enzyme 1 and 5 μl of Enzyme 2 (Pierce Cardiomyocytes Dissociation Kit, ThermoFisher Scientific) for 30 min at 350 rpm shaking at 30°C. The cells were centrifuged for 5 min at 5000 rpm, and the dissociation media was removed and replaced by DMEM+10% FBS. The cells were then sorted using a BD FACSAria^TM^ III – firstly by forward scatter amplitude and side scatter amplitude, and subsequently sorted by GFP and mCherry fluorescence into Trizol. RNA extraction was then performed using Qiagen RNeasy Kit. At least 100 ng of total RNA was used for reverse transcription using Maxima First Strand cDNA synthesis kit (Thermo Fisher Scientific). DyNAmo ColorFlash SYBR Green qPCR Mix (Thermo Fisher Scientific) was used on a CFX connect Real-time System (Bio-Rad) with the following program: pre-amplification 95°C for 7 min, amplification 95°C for 10 s and 60°C for 30 s (repeated for 39 cycles), melting curve 60-92°C with increment of 1°C each 5 s. Each point in the dot plots represents a biological replicate from three technical replicates. Gene expression values were normalized using the housekeeping gene *rpl13a* and fold changes were calculated using the 2^-ΔΔCt^ method.

### Zebrafish transgenesis

Tol2 transgenesis was used to generate the *Tg(fli1:gpr126-p2a-tdTomato)* and *Tg(fli1:tdTomato-p2a-gpr126CTF)* lines. To generate stable lines, wild-type AB embryos (F0) were injected at the one-cell stage with an injection cocktail of transposase (tol2 mRNA, 12.5 pg per embryo) and plasmid DNA (25 pg per embryo). F0 embryos positive for tdTomato fluorescence were raised to adulthood and then screened for founder animals. The founders (three for each transgenic line) were further outcrossed to raise the F1 generation.

### Morpholino injection

In order to stop contractility and blood flow in the embryos, 0.25 ng of *tnnt2a* morpholino (5′-CATGTTTGCTCTGATCTGACACGCA-3′) was injected into single cell-stage embryos.

### Chemical treatments

ErbB2 signaling was inhibited by treating the embryos with 10 μM of ErbB2 inhibitor PD168393 (Calbiochem).10 µM working solution of PD168393 was prepared in 1x E3 media with 0.003% PTU. For the treatment, 10 embryos per well were placed in a 6-well plate containing the working solution of PD168393. As a control 1% DMSO was used. The treatment was performed from 60 hpf to 120 hpf. At 120 hpf, the PD168393 solution was removed, and the embryos were washed three times with 1x E3 medium for 5 min and subsequently imaged.

### Imaging

Image acquisition for immunostained cells as well as zebrafish hearts was performed using Carl Zeiss AG LSM800 confocal laser scanning microscope equipped with the ZEISS blue software. For live imaging of stopped zebrafish hearts, embryos and larvae were mounted in 1% low melting agarose on glass-bottom dishes (MatTek). The larvae were oriented through a micromanipulator under the Carl Zeiss AG Axiozoom V16 stereoscope to enhance optical access to the heart. In order to stop the heart beating before imaging, 0.2% (w/v) tricaine was added to the low melting agarose. A 40x (1.1 numerical aperture [NA]) water-immersion objective was used to acquire the images. The optical sections were 1 µm thick.

### Image processing and Quantification

2D and 3D images were processed using ImageJ and the Fiji extension package. Figures were prepared with the help of Adobe Illustrator or Microsoft PowerPoint. The quantification of intensity profiles of HA- and GFP-tag in ARPE-19 cells were performed using the “Measure” tool. The GFP intensity for each condition was measured relative to the HA intensity. For the quantification of myocardial Notch activity, intensity profiling of the YFP signal was performed. The total YFP signal relative to the YFP signal in the valve endocardium was plotted as intensity of myocardial Notch. Three-dimensional surface rendering was performed using the Surfaces function of Imaris x64 (Bitplane). Bright-field images obtained from the stereoscope were processed using Zen blue (ZEISS) software.

### Statistical Analysis

Statistical analysis and p-value (p) calculation was performed using Graphpad Prism 8.3.0. Data are expressed as mean ± SEM or S.D as indicated. Differences between two groups were determined with a two-tailed Student’s t-distribution test. To compare multiple conditions, one-way ANOVA was performed and when data were compared to control groups, correction for multiple comparisons was performed by using Dunnett’s test. All tests were performed with a confidence level of 95%.

## Funding

This work was supported by the Deutsche Forschungsgemeinschaft (DFG, INST 410/91-1 FUGG as well as FOR2149, Project 7, EN 453/10-1 and EN 453/10-2 to F.B.E and Bavarian Equal Opportunities Sponsorship – Realisierung von Frauen in Forschung und Lehre (FFL) – Promotion Equal Opportunities for Women in Research and Teaching funding to S.S. The building of *stl47* allele was supported by NIH NINDS F32 NS087786 funding to S.C.P.

## Acknowledgements

We are grateful to Kelly Monk, Robert Becker, and Silvia Vergarajauregui for discussions and critical reading of the manuscript. We would like to thank Ines Liebscher for zebrafish Gpr126 constructs, Sebastian Zundler for providing IMARIS software, Darren Gilmour for providing *TgBAC(cdh2:cdh2-GFP)*, as well as Jana Petzold, Christina Goula, and Jennifer Redlingshöfer for excellent care of the zebrafish facility. SS acknowledges the support from Filomena Ricciardi for sharing her expertise in trabeculation imaging.

## Author Contribution

S.S. designed, performed and analyzed most of the experiments. F.G and A.G performed the FACS sorting for endocardial and myocardial cells from zebrafish larval hearts and provided *gpr126* expression data in the sorted cells. S.C.P generated and provided the *stl47* allele. S.S and F.B.E. designed all experiments, interpreted results, and wrote the manuscript. D.Y.R.S provided constant guidance with the project as well as various transgenic lines. F.B.E supervised the project. All the authors contributed to the discussion of data and refining of the manuscript.

## Declaration of Interests

The authors have no conflict of interest to declare.

---

**Figure S1.**
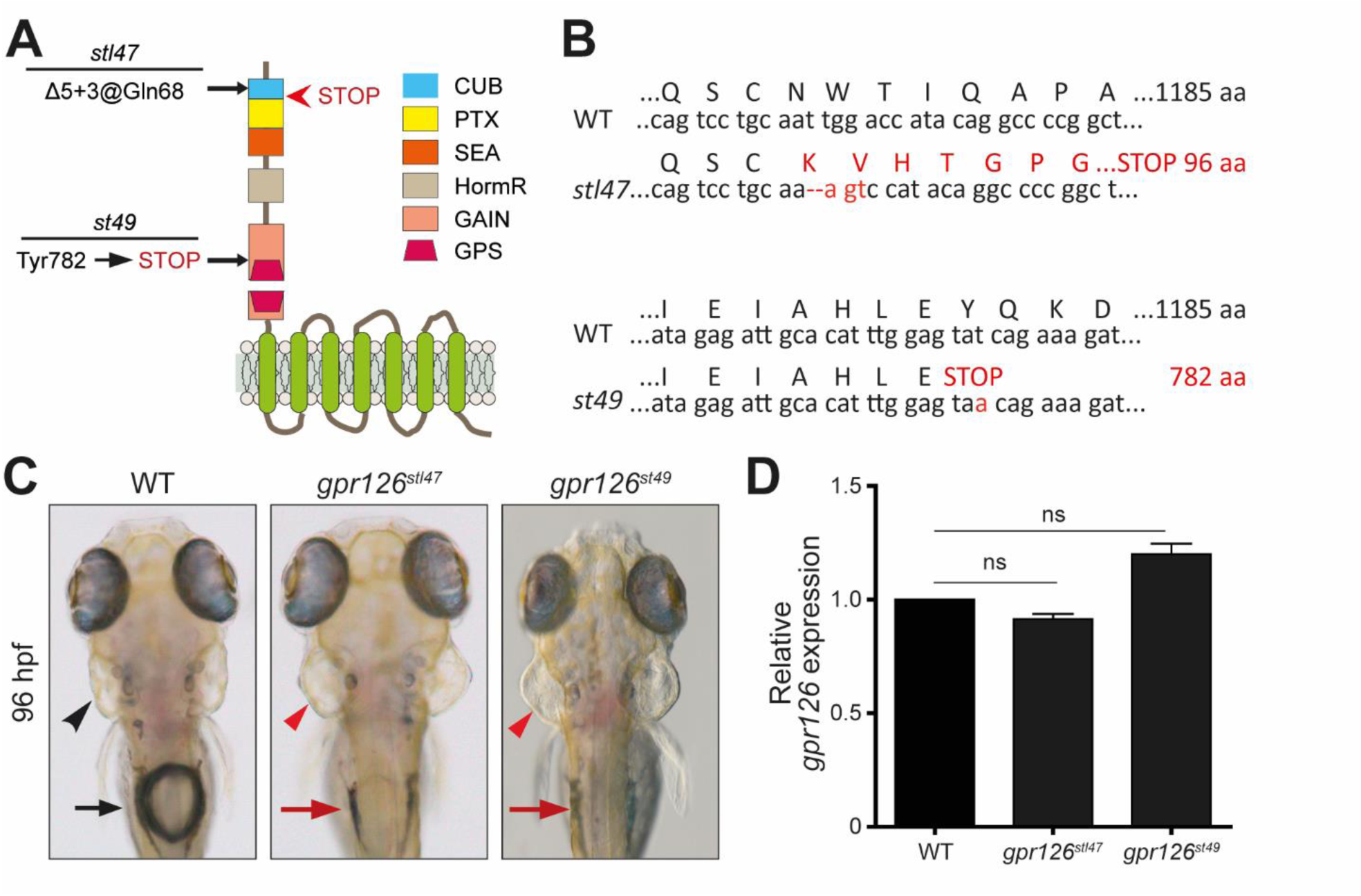
*gpr126^stl47^* and *gpr126^st49^* mutants exhibit no nonsense-mediated RNA decay. **(A)** Schematic representation of zebrafish Gpr126 indicating changes encoded by *stl47* and *st49* alleles. The red arrowhead indicates the STOP codon encoded by *stl47* allele. **(B)** Sequence comparison of WT vs. *stl47* allele and WT vs. *st49* allele. WT gpr126 encodes for an 1185 aa protein. The *stl47* allele has a Δ5+3 indel that should result in truncation of the protein at aa 96. The *st49* allele carries a point mutation that converts a tyrosine-encoding codon into a stop codon resulting in a truncation of the protein at aa 782. **(C)** Dorsal views of WT, *gpr126^stl47^*, and *gpr126^st49^* larvae at 96 hpf. Mutants exhibit puffy ear and swim bladder inflation defects. Black arrowhead: normal ear; black arrow: inflated swim bladder; red arrowhead: puffy ear; red arrow: failure of swim bladder to inflate. **(D)** qPCR expression data of *gpr126* in 96 hpf hearts from WT, *gpr126^stl47^*, and *gpr126^st49^* larvae. Data are mean ± SD; n = 3; ns: not statistically significant.

**Figure S2.**
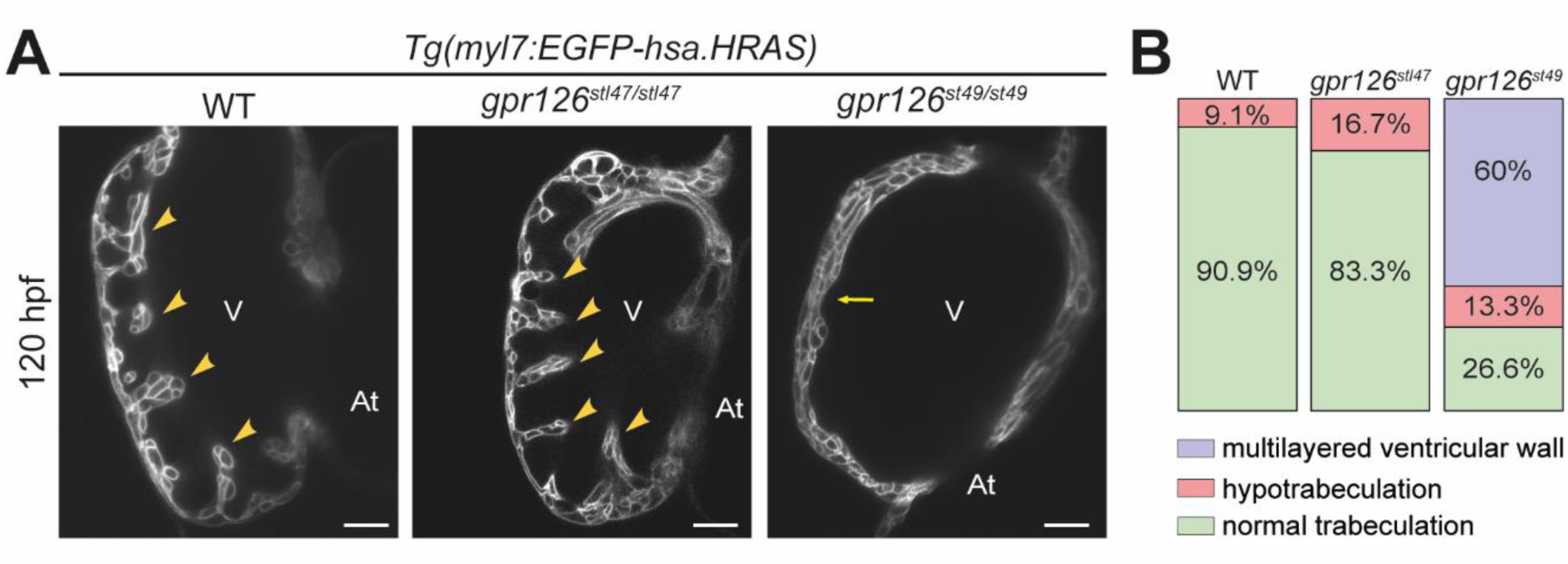
Zygotic *gpr126^stl47^* hearts do not exhibit trabeculation defects. **(A)** Representative confocal images (mid-sagittal sections) of hearts of WT, *gpr126^stl47/stl47^*, and *gpr126^st49/st49^* larvae crossed into *Tg(myl7:EGFP-hsa.HRAS)* background at 120 hpf. Arrowheads: trabeculae, Arrows: multilayered ventricular wall. V: ventricle. At: atrium. Scale bars: 20 µm. **(B)** Quantification of trabeculation phenotypes in 120 hpf WT (n=11), *gpr126^stl47/stl47^* (n=12), and *gpr126^st49/st49^* (n=15) larvae.

**Figure S3.**
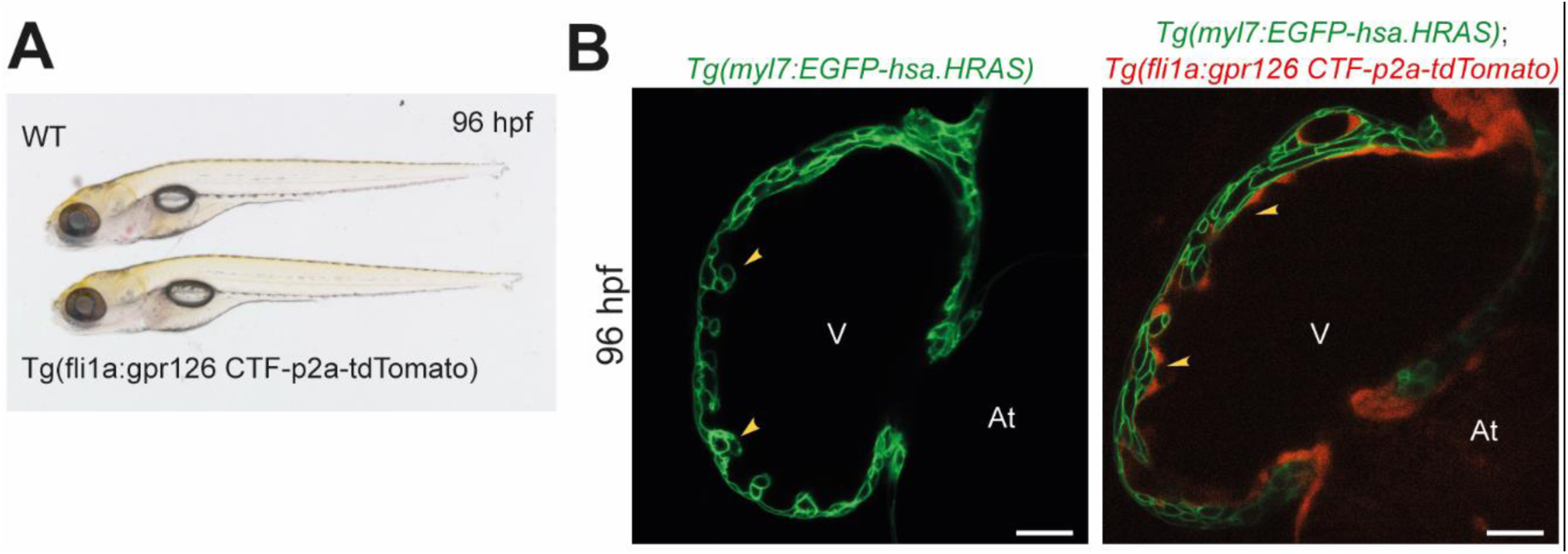
Endocardial CTF overexpression leads to no obvious heart defects. **(A)** Representative gross morphology of 96 hpf WT and *Tg(fli1a:gpr126 CTF-p2a-tdTomato)* larvae. **(B)** Confocal images (mid-sagittal sections) of *Tg(my7:EGFP-hsa.HRAS)* and *Tg(my7:EGFP-hsa.HRAS;Tg(fli1a:gpr126 CTF-p2a-tdTomato)* at 96 hpf. Arrowheads: trabeculae; V: Ventricle; At: Atrium; Scale bars: 20 µm.

**Figure S4.**
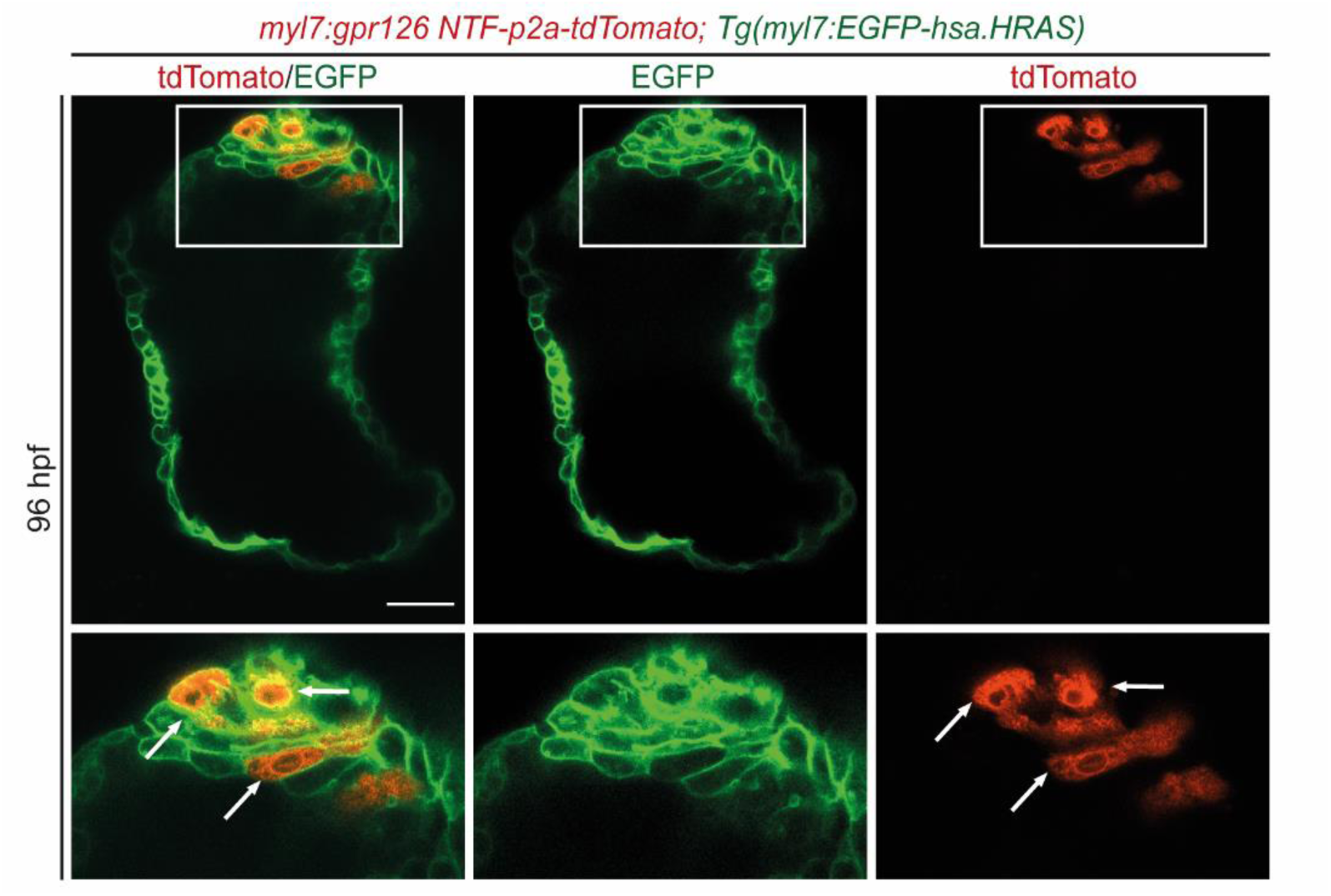
Mosaic myocardial-specific NTF overexpression induces local cardiomyocyte multilayering. Confocal images (mid-sagittal sections) of 96 hpf *Tg(myl7:EGFP-has.HRAS*) hearts injected with a myocardial-specific NTF overexpression construct *(myl7:gpr126 NTF-p2a-tdTomato)* at the single cell-stage. Mosaic overexpression of NTF in WT cardiomyocytes leads to local multilayering (5/8) which is outlined by a white box and magnified. Arrows: NTF overexpressing cardiomyocytes. Scale bars: 20 µm.

**Figure S5.**
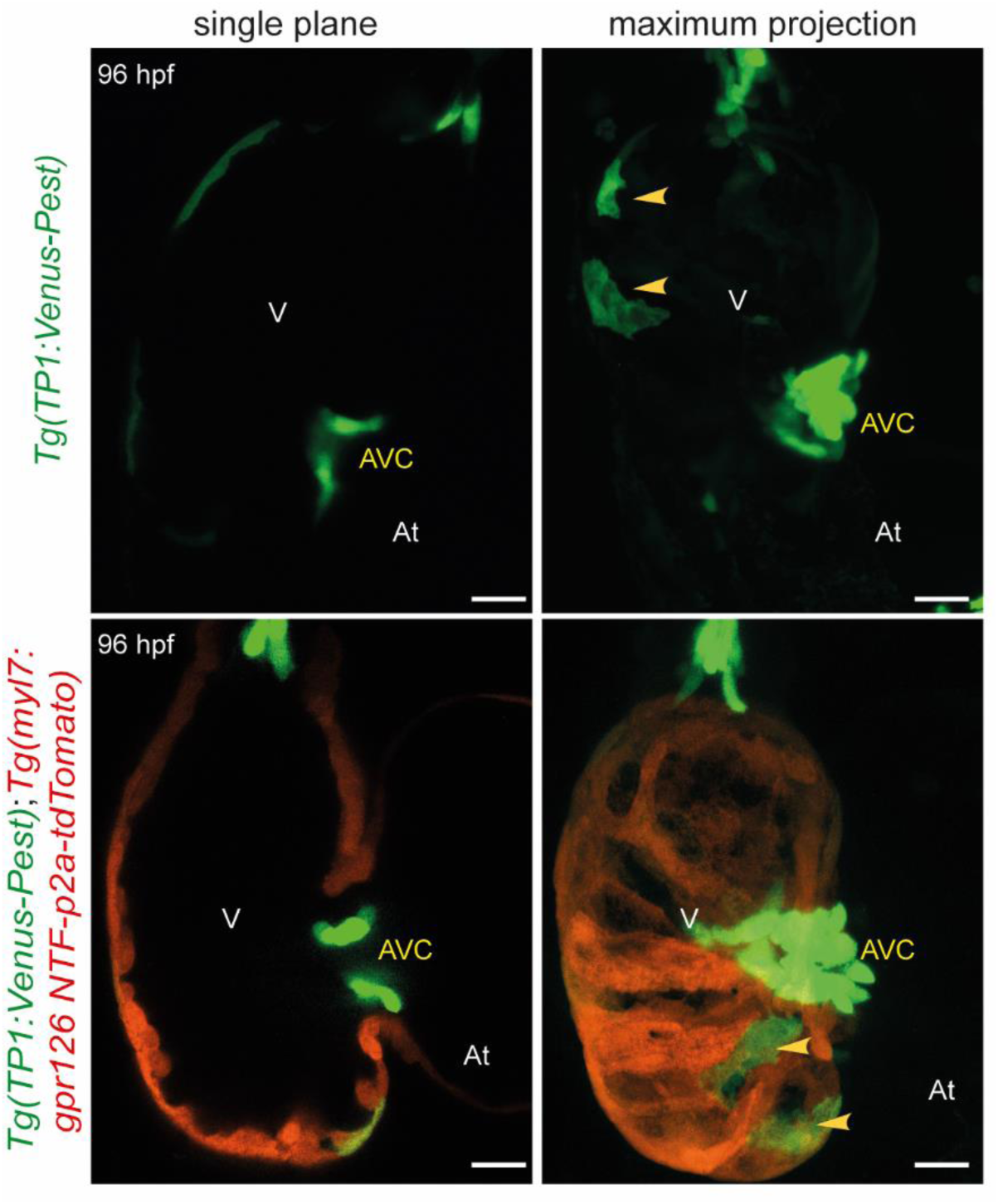
NTF overexpression in the myocardium does not lead to an increase in myocardial Notch activity. Confocal images of 96 hpf *Tg(TP1:Venus-Pest*) and *Tg(Tp1:Venus-Pest); Tg(myl7:gpr126 NTF-p2a-tdTomato)* hearts. Arrows: myocardial Notch activity; Scale bars: 20 µm.

## Notes

### Competing Interest Statement

The authors have declared no competing interest.

